## Supplemental figures and tables for "Multi-omic characterization of allele-specific regulatory variation in hybrid pigs": Figure S4.pdf

| F40 | F70 | D1 | D168 | Brain | Liver | Muscle | Placenta (F40 vs. F70) |
|---|---|---|---|---|---|---|---|
| ○ | → | → | → | 714 | 403 | 301 | 9832 |
| ○ | → | → | ↑ | 283 | 404 | 431 | — |
| ○ | → | → | ↓ | 236 | 291 | 462 | — |
| ○ | → | ↑ | → | 809 | 444 | 563 | — |
| ○ | → | ↑ | ↑ | 564 | 478 | 755 | — |
| ○ | → | ↑ | ↓ | 199 | 630 | 955 | — |
| ○ | → | ↓ | → | 921 | 545 | 579 | — |
| ○ | → | ↓ | ↑ | 210 | 606 | 808 | — |
| ○ | → | ↓ | ↓ | 544 | 367 | 917 | — |
| ○ | ↑ | → | → | 679 | 310 | 128 | 1269 |
| ○ | ↑ | → | ↑ | 263 | 472 | 212 | — |
| ○ | ↑ | → | ↓ | 228 | 222 | 268 | — |
| ○ | ↑ | ↑ | → | 1128 | 509 | 287 | — |
| ○ | ↑ | ↑ | ↑ | 739 | 807 | 574 | — |
| ○ | ↑ | ↑ | ↓ | 287 | 528 | 546 | — |
| ○ | ↑ | ↓ | → | 617 | 256 | 307 | — |
| ○ | ↑ | ↓ | ↑ | 167 | 523 | 425 | — |
| ○ | ↑ | ↓ | ↓ | 508 | 222 | 527 | — |
| ○ | ↓ | → | → | 727 | 309 | 200 | 1122 |
| ○ | ↓ | → | ↑ | 230 | 183 | 330 | — |
| ○ | ↓ | → | ↓ | 233 | 397 | 203 | — |
| ○ | ↓ | ↑ | → | 601 | 313 | 369 | — |
| ○ | ↓ | ↑ | ↑ | 348 | 230 | 458 | — |
| ○ | ↓ | ↑ | ↓ | 188 | 584 | 522 | — |
| ○ | ↓ | ↓ | → | 1201 | 625 | 515 | — |
| ○ | ↓ | ↓ | ↑ | 235 | 549 | 704 | — |
| ○ | ↓ | ↓ | ↓ | 917 | 880 | 539 | — |
