## Supplementary figures and images for "Multi-omic characterization of allele-specific regulatory variation in hybrid pigs"

### Figure S1.jpg

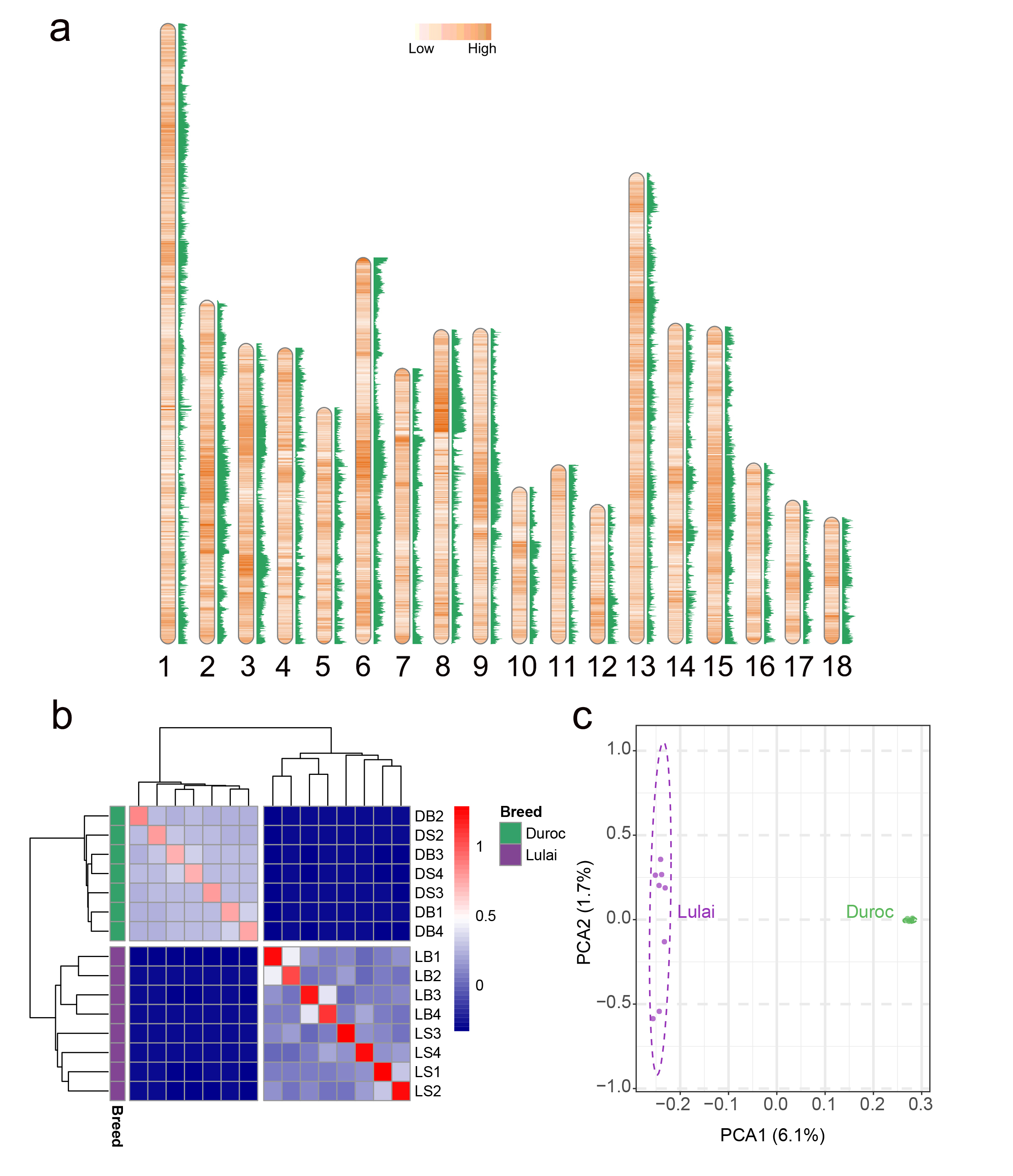

### Figure S2.pdf

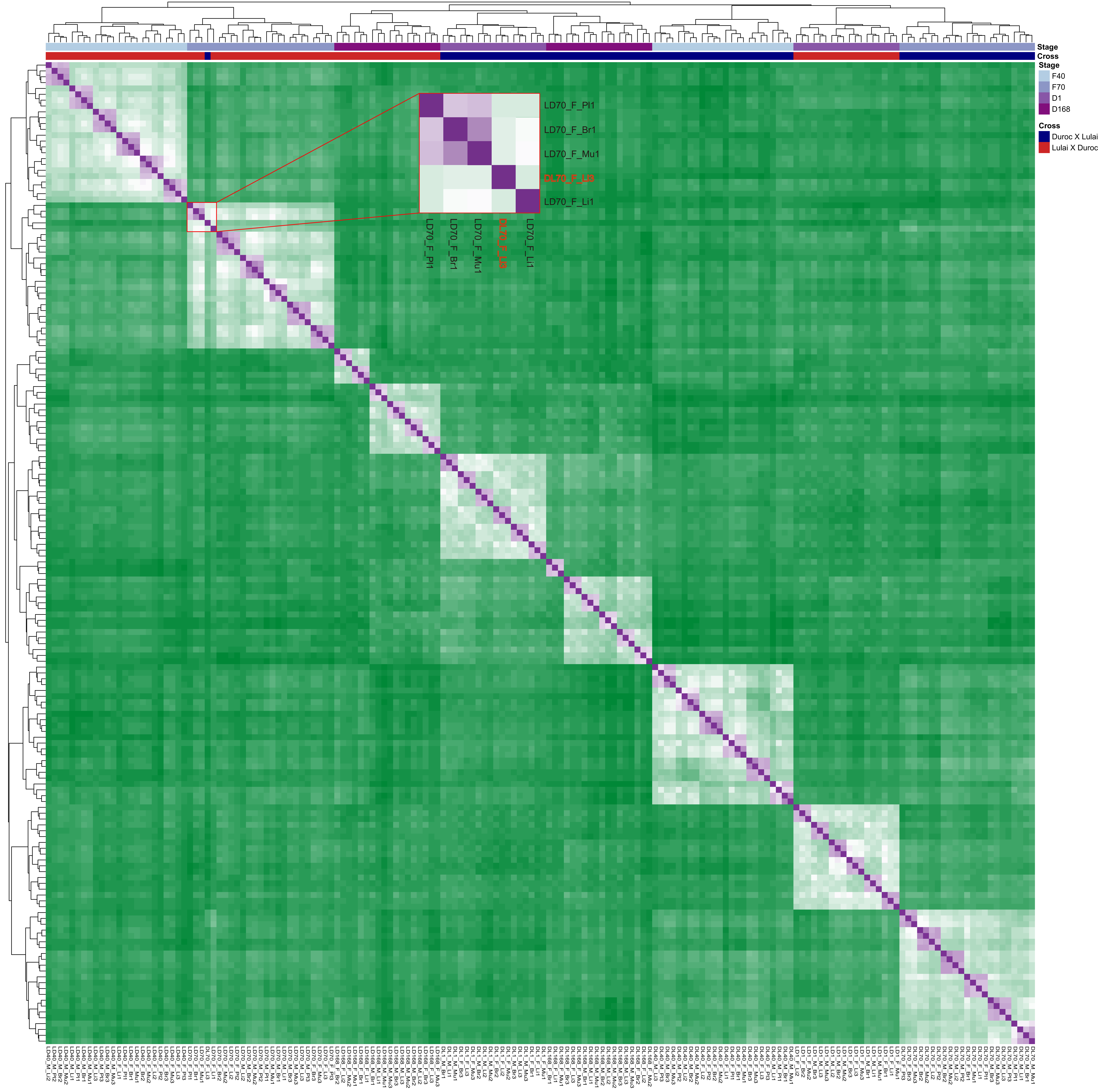

### Figure S3.png

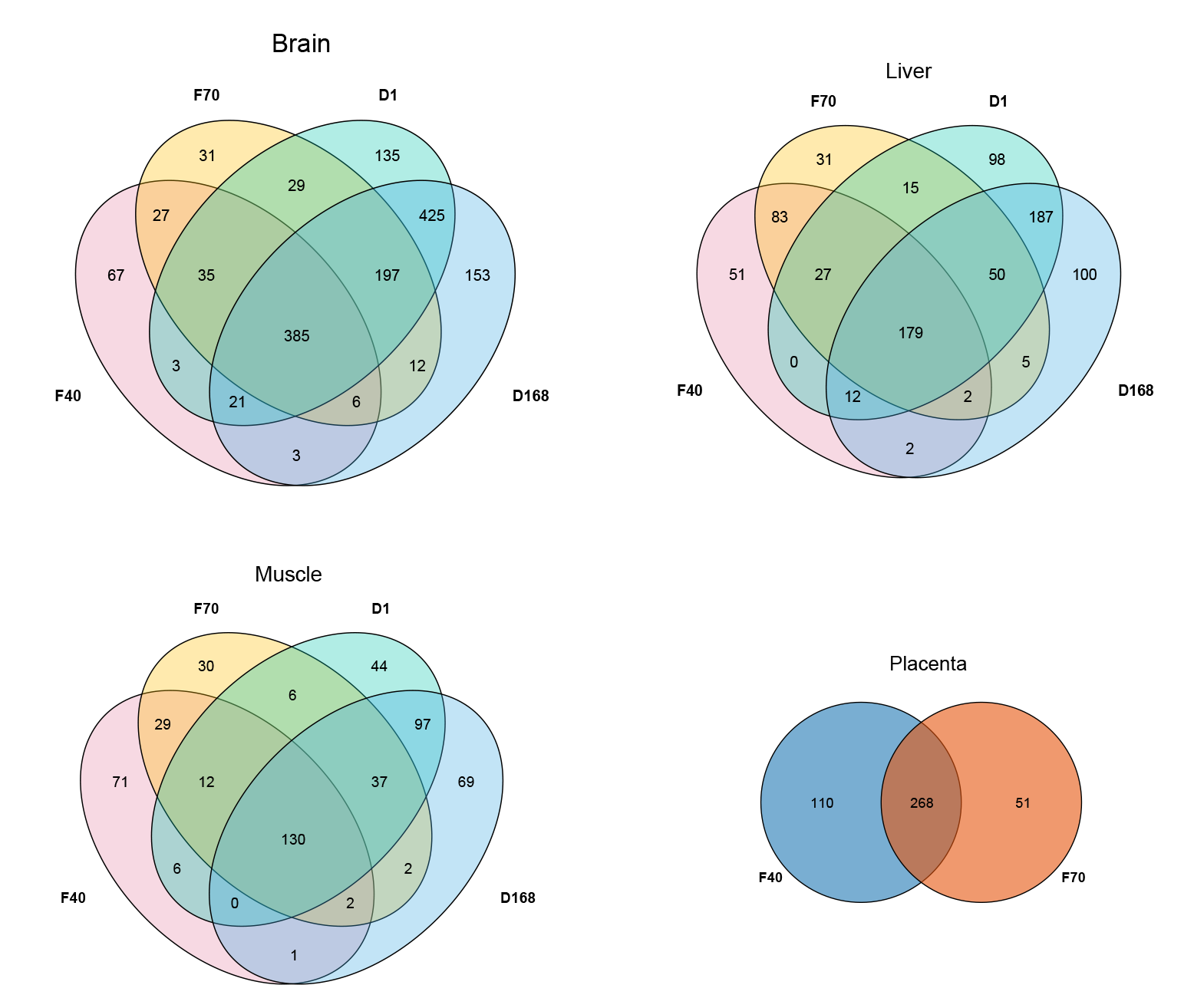

### Figure S5.pdf

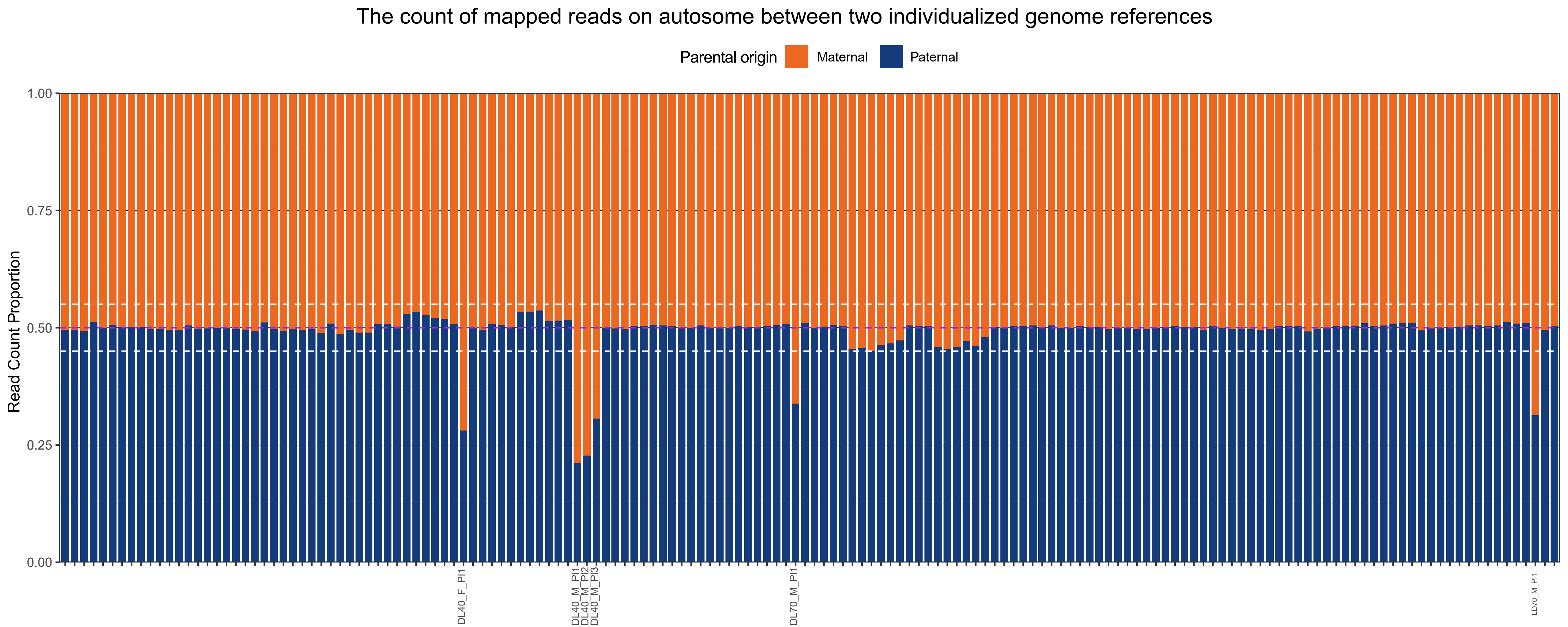

### Figure S6.jpg

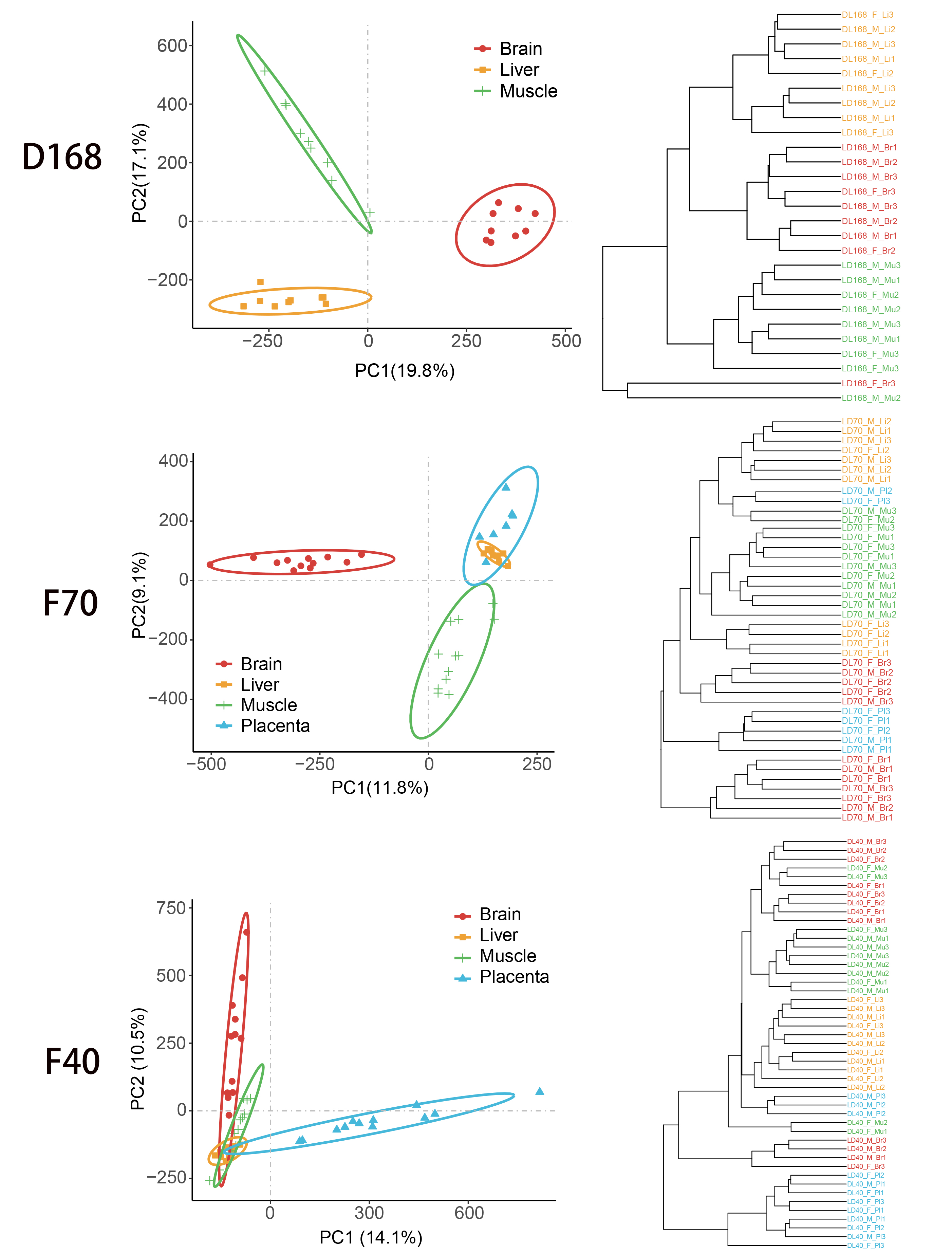

### Figure S7.pdf

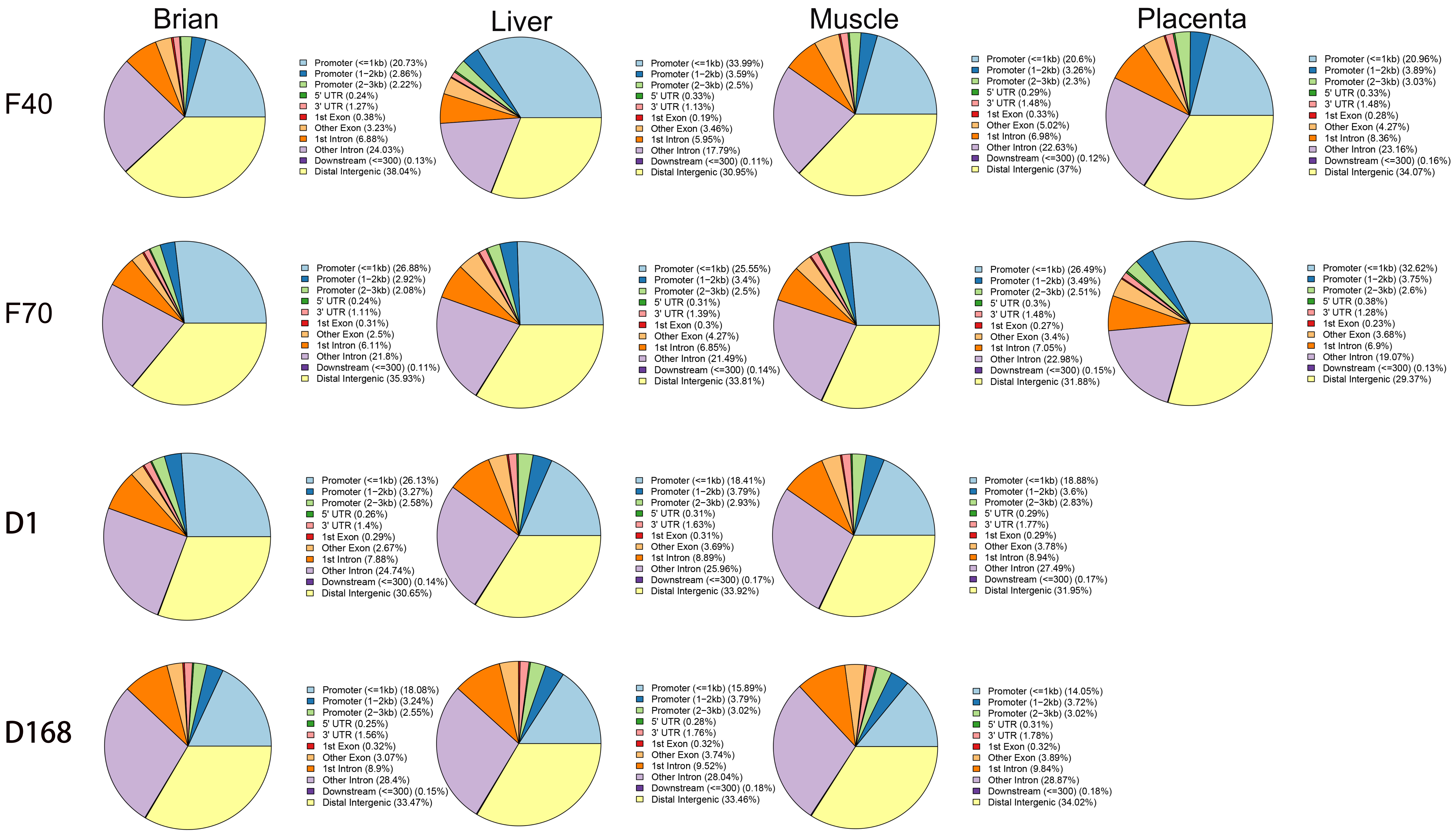

### Figure S8.pdf

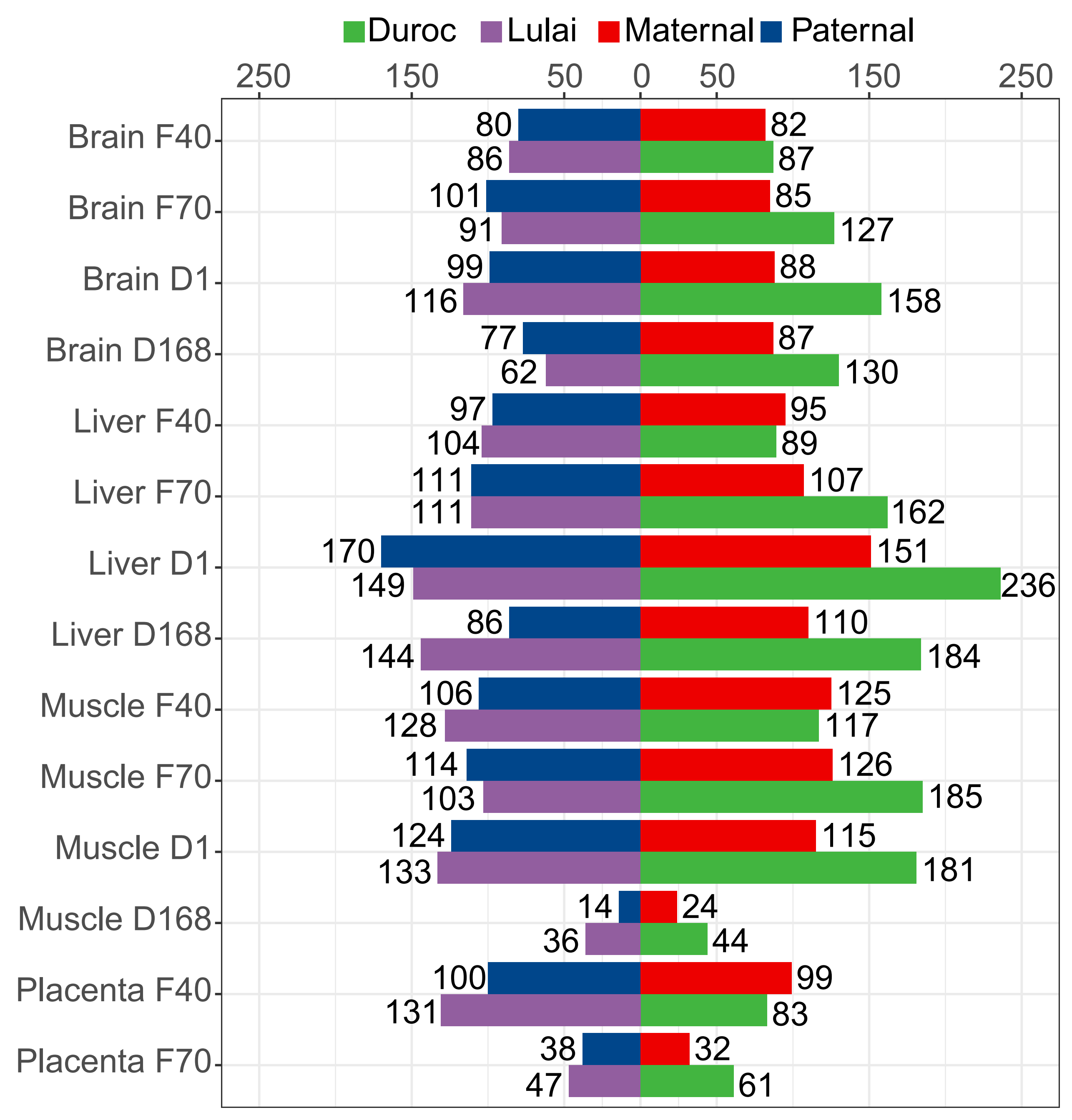

### Figure S9.pdf

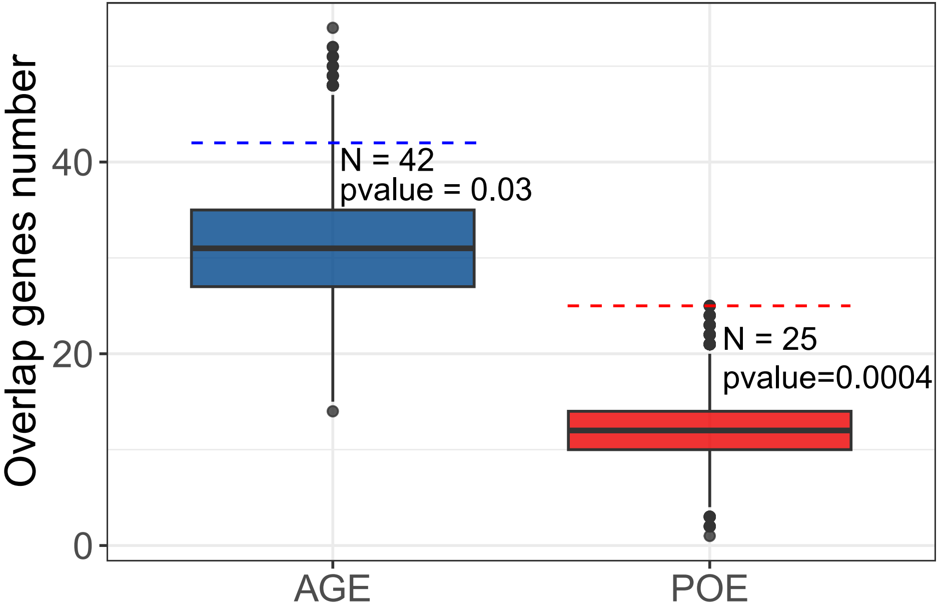

### Figure S10.jpg

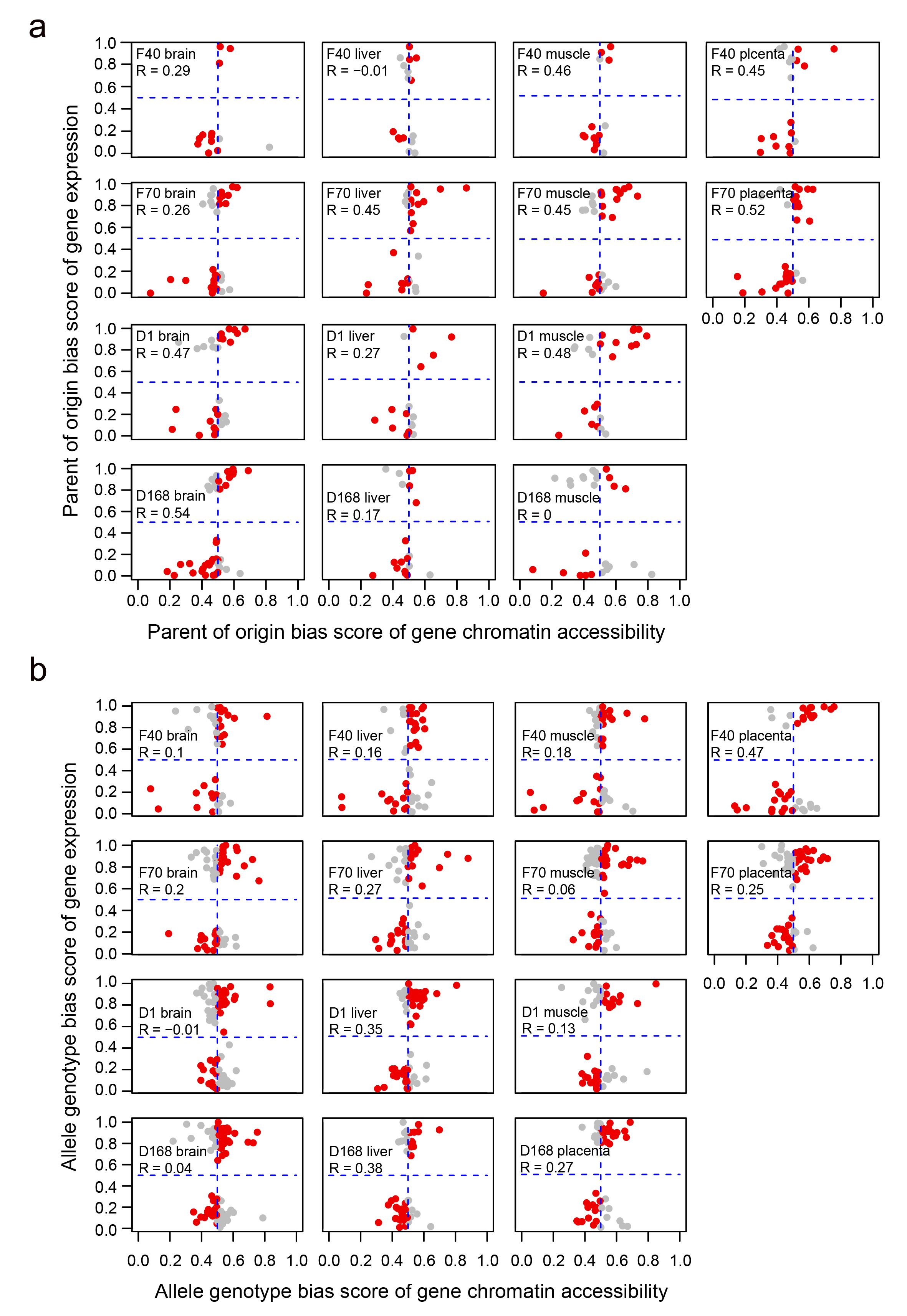

### Figure S11.pdf

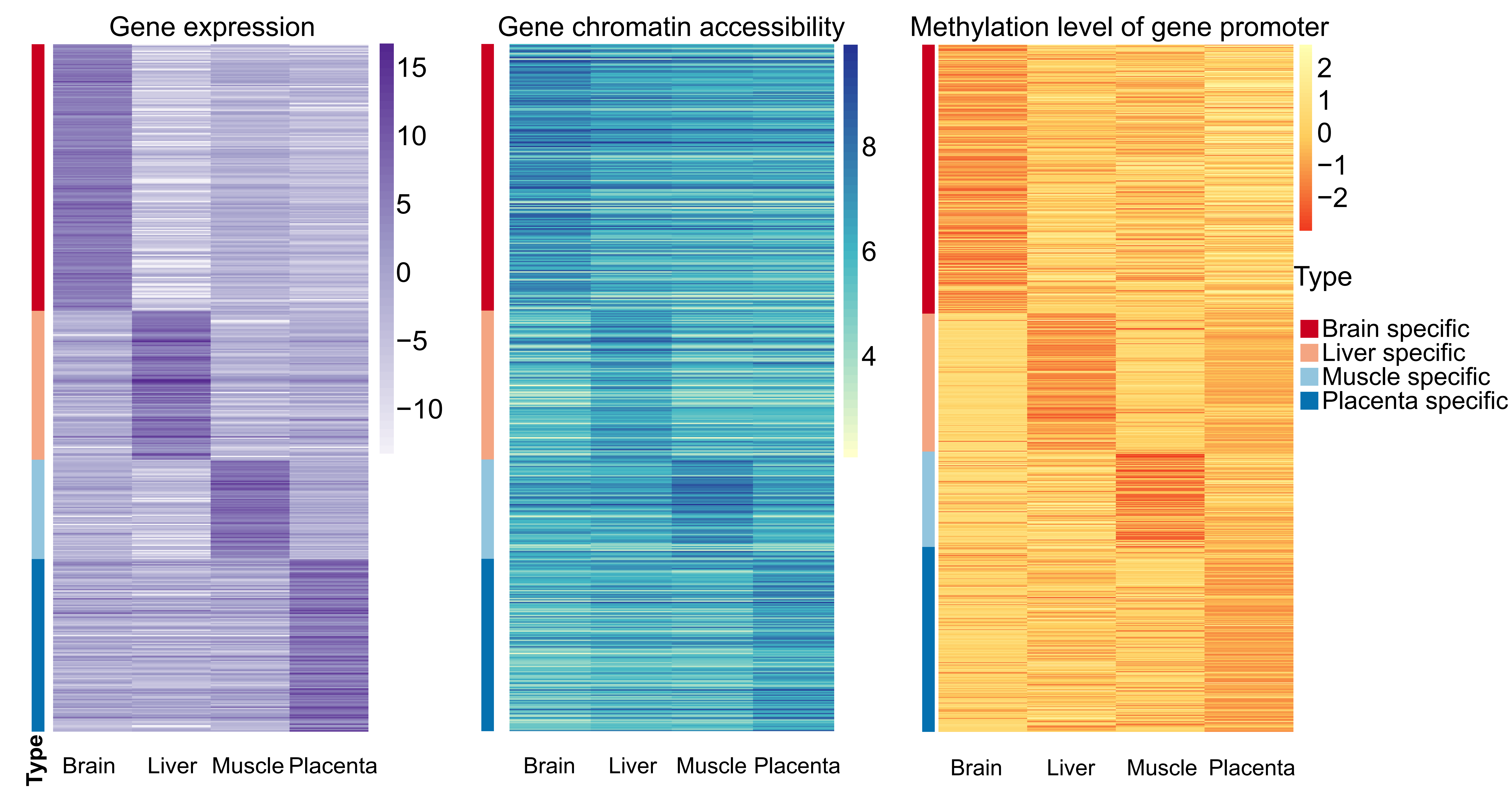

### Figure S12.pdf

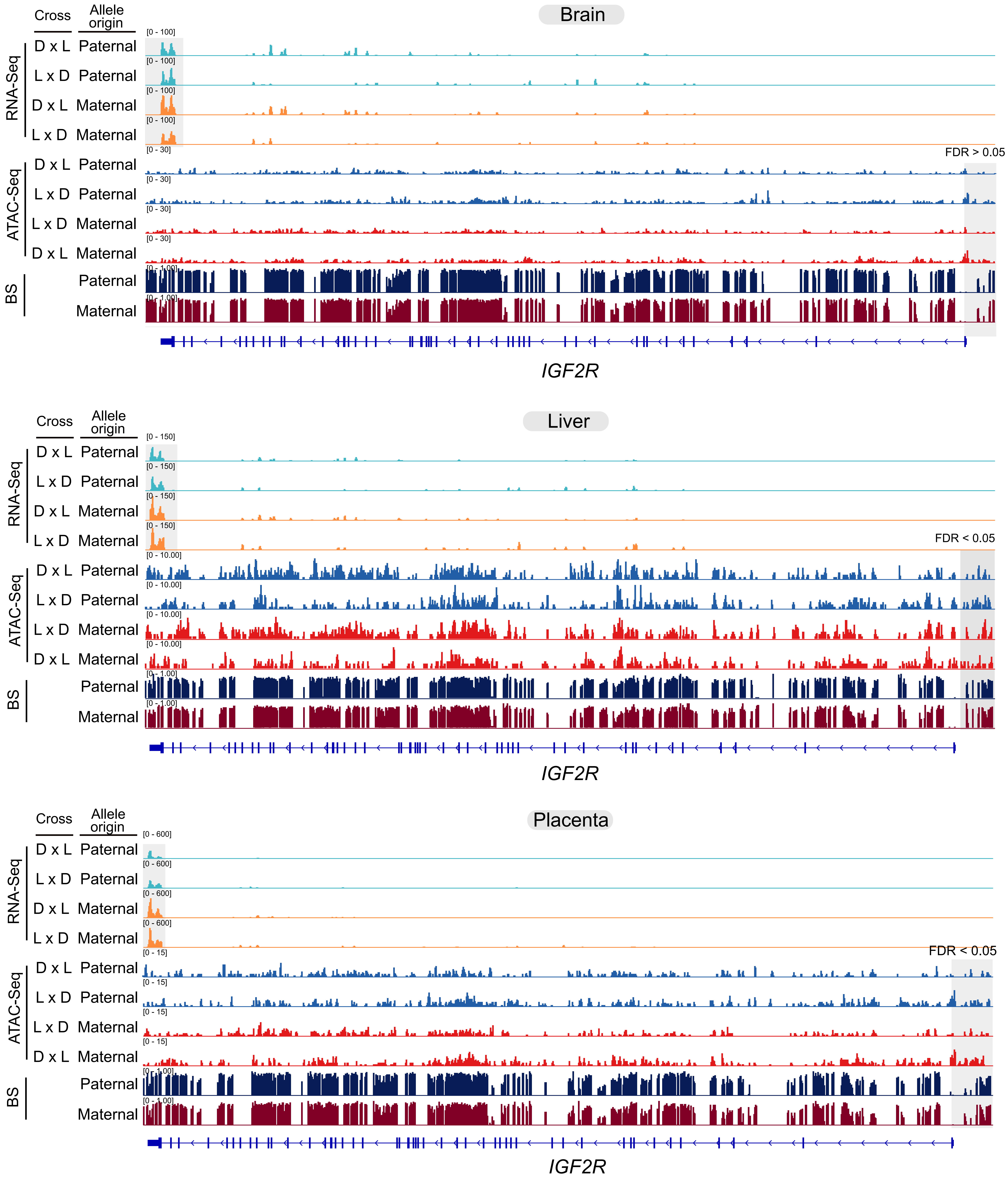

### Figure S13.pdf

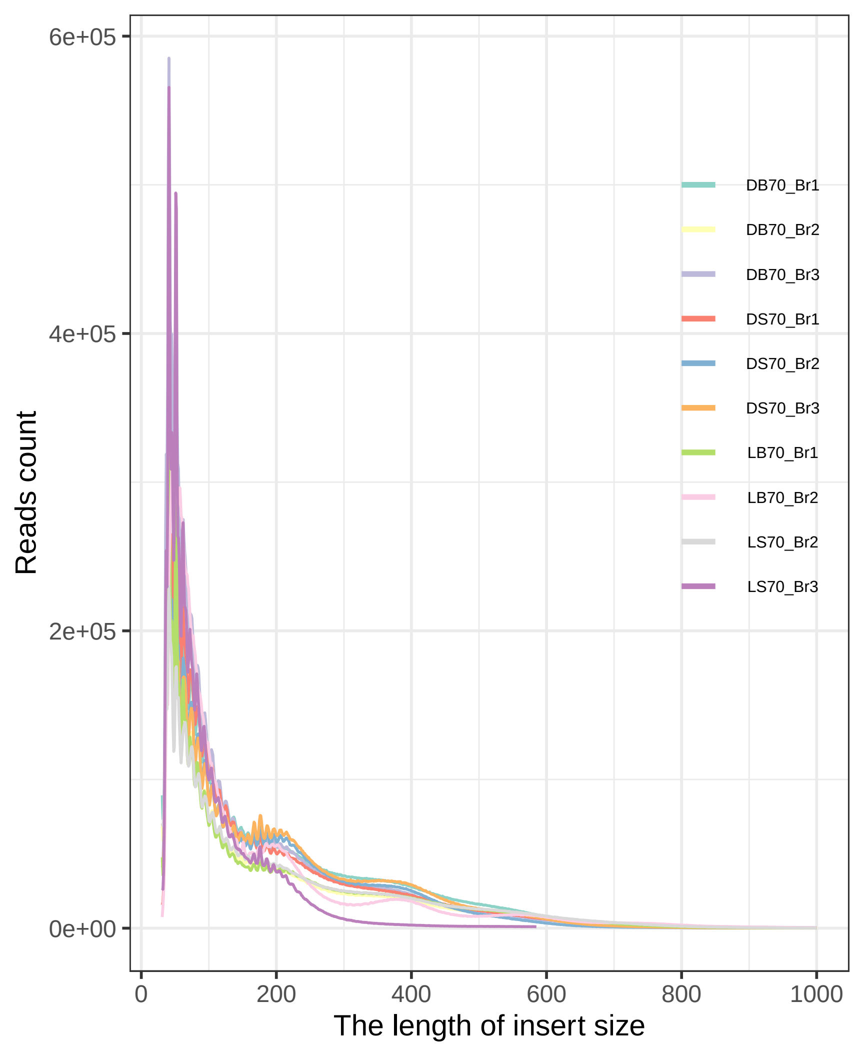
